# PAH DEFICIENT PATHOLOGY IN HUMANIZED c.1066-11G>A PHENYLKETONURIA MICE

**DOI:** 10.1101/2023.11.03.565447

**Authors:** Ainhoa Martínez-Pizarro, Sara Picó, Arístides López-Márquez, Claudia Rodriguez-López, Elena Montalvo, Mar Alvarez, Margarita Castro, Santiago Ramón-Maiques, Belén Pérez, José J Lucas, Eva Richard, Lourdes R Desviat

**Author notes:** Correspondence should be addressed to: Lourdes R Desviat, Centro de Biología Molecular Severo Ochoa, Universidad Autónoma de Madrid Nicolás Cabrera 1, 28049 Madrid, Spain. Department of Biomedical Sciences, Faculty of Biology and Medicine, University of Lausanne, Lausanne, Vaud, Switzerland (Sara Picó). Institut de Recerca de Sant Joan de Déu, Barcelona, Spain (Arístides López-Márquez). The authors have declared that no conflict of interest exists.

## Abstract

We have generated using CRISPR/Cas9 technology a partially humanized mouse model of the neurometabolic disease phenylketonuria (PKU), carrying the highly prevalent *PAH* variant c.1066-11G>A. This variant creates an alternative 3’ splice site, leading to the inclusion of 9 nucleotides coding for 3 extra amino acids between Q355 and Y356 of the protein. Homozygous *Pah* c.1066-11A mice, with a partially humanized intron 10 sequence with the variant, accurately recapitulate the splicing defect and present almost undetectable hepatic PAH activity. They exhibit fur hypopigmentation, lower brain and body weight and reduced survival. Blood and brain phenylalanine levels are elevated, along with decreased tyrosine, tryptophan and monoamine neurotransmitter levels. They present behavioral deficits, mainly hypoactivity and diminished social interaction, locomotor deficiencies and an abnormal hind-limb clasping reflex. Changes in the morphology of glial cells, increased GFAP and Iba1 staining signals and decreased myelinization are observed. Hepatic tissue exhibits nearly absent PAH protein, reduced levels of chaperones DNAJC12 and HSP70 and increased autophagy markers LAMP1 and LC3BII, suggesting possible coaggregation of mutant PAH with chaperones and subsequent autophagy processing. This PKU mouse model with a prevalent human variant represents a useful tool for pathophysiology research and for novel therapies development.

## Introduction

Phenylketonuria (PKU; OMIM#261600) is probably the best studied and the most frequent inborn error of amino acid metabolism, caused by autosomal recessive deficiency of the hepatic enzyme phenylalanine hydroxylase (PAH; EC 1.14.16.1), catalysing the (6R)-5,6,7,8-tetrahydrobiopterin (BH_4_) dependent hydroxylation of L-phenylalanine (Phe) to L-tyrosine (Tyr). The global PKU prevalence was recently estimated to be approximately 1:24,000 newborns (1), with a documented large frequency variability depending on the population and geographical location, being around 1:10,000 in USA and parts of Europe (1). In PAH deficiency, high Phe levels cause brain dysfunction in untreated patients, exhibiting intellectual disability, seizures and behavioural problems, although a broad clinical spectrum exists, from severe, classical PKU to the mildest form, mild hyperphenylalaninemia (HPA) with normal brain development (2). Early diagnosis is based on newborn screening and implementation of Phe-restricted dietary treatment, which has been the gold standard for many decades (3). This prevents major neurological alterations, although significant challenges remain, associated to poor adherence and suboptimal outcomes. In the last years, new treatment options have become available, a synthetic form of BH_4_ (sapropterin, Kuvan®) which allows a less restrictive or absence of diet for a group of patients with mild PKU forms, and a pegylated form of phenylalanine ammonia lyase, indicated for adult patients (2). Other treatments are in preclinical or clinical development, such as mRNA, gene and gene editing therapies, or use of a probiotic *Escherichia coli* Nissle strain, engineered to overexpress enzymes metabolizing gut Phe (4).

In order to investigate PKU pathophysiology and as tools for preclinical novel treatment testing, several mouse models have been generated. First models were obtained by N-ethyl-N-nitrosourea germline mutagenesis, with *Enu1* mice representing a mild HPA phenotype and *Enu2* and *Enu3* as models for severe, classical PKU (5, 6). However, the murine mutations present in these models do not correspond to human variants, and, with the advent of gene editing technologies, mouse “avatar” models carrying variants identified in human patients (p.R261Q, p.R408Wy, p.P281L) have now been generated (7–10). To date, no mouse models with *PAH* splicing variants have been characterised, despite their frequency in the PKU mutational spectrum. The second most frequent variant worldwide is c.1066-11G>A (global allele frequency 6.4%) (1), which is often called the ‘‘Mediterranean mutation,’’ as it accounts for the majority of the mutant alleles in Southern Europe (11%), and is also common in Latin America (9,4%). The homozygous genotype c.[1066-11G>A];[1066-11G>A] is the second most prevalent genotype worldwide (2,6% globally) and the first in Southern Europe (5,2%) (1), associated to a classical, non BH_4_-responsive severe phenotype (www.biopku.org).

The *PAH* c.1066-11G>A variant, located in intron 10, creates a strong splice acceptor site used instead of the natural one, leading to the aberrant in-frame insertion of 9 nucleotides in the mature mRNA, coding for three extra amino acids between residues 355 and 356 of the protein (p.Q355_Y356insGLQ). The initial study reported the presence of immunoreactive protein in a liver biopsy from a homozygous patient but total absence of catalytic activity (11). Expression analysis in a prokaryotic system of the mutant protein (with the 3 extra amino acids) fused to maltose binding protein showed that it was recovered exclusively as high molecular weight aggregates, with no catalytic activity (12). The exact liver pathomechanism associated to the mutant protein is yet unknown, stressing the need of finding an adequate *in vivo* model in which to test potential therapies.

The fact that intronic sequences are not conserved between mouse and human, and that, even with an equivalent nucleotide change, the splicing outcome may be different due to the distinct genomic context of the mutation (13), hinders the precise modelling of intronic splicing defects in mice. To that end, based on the interest in the prevalent c.1066-11G>A variant, in this work we have used CRISPR/Cas9 technology to generate a PKU mouse model with a partially humanized *Pah* intron 10 carrying the variant. After confirming the altered *Pah* splicing profile in liver of these so called *Pah* c.1066-11A *knock in* (*KI*) mice, accurately recapitulating the human molecular defect, we have characterised their biochemical, histological, and behavioural phenotypes. We have also used molecular approaches to investigate pathological mechanisms underlying mutant protein expression in liver. In parallel, we have generated a stable isogenic cellular model of the splicing variant through CRISPR/Cas gene editing. To this end, since *PAH* is expressed mainly in liver, we decided to use HepG2 cells as the most suitable cell line to study the characteristics of the mutant protein in a human genetic background. The mouse avatar model and the edited HepG2 cell line, both carrying the c.1066-11A variant corresponding to a frequent classical PKU phenotype, are ideally suited for preclinical research and development of RNA therapies targeting the splicing defect or pharmacological compounds targeting the mutant protein.

## Results

### The c.1066-11A *PAH* variant minigene mimics disease-associated mis-splicing

Bioinformatic analysis of *PAH* intron 10 sequence confirmed the creation of a 3’ splice acceptor site with variant c.1066-11G>A (MaxEnt score 5.5), which is predicted to be used instead of the natural splice site (MaxEnt score 3.2). Functional analysis in the pcDNA3.1 minigene (14) confirmed the splicing defect (Figure 1A). As previously reported, residual exon 11 skipping is observed in the wild type (WT) minigene, due to a naturally weak 3’ splice site and a suboptimal 5’ splice site (14) (Figure 1A).

**Figure 1.**
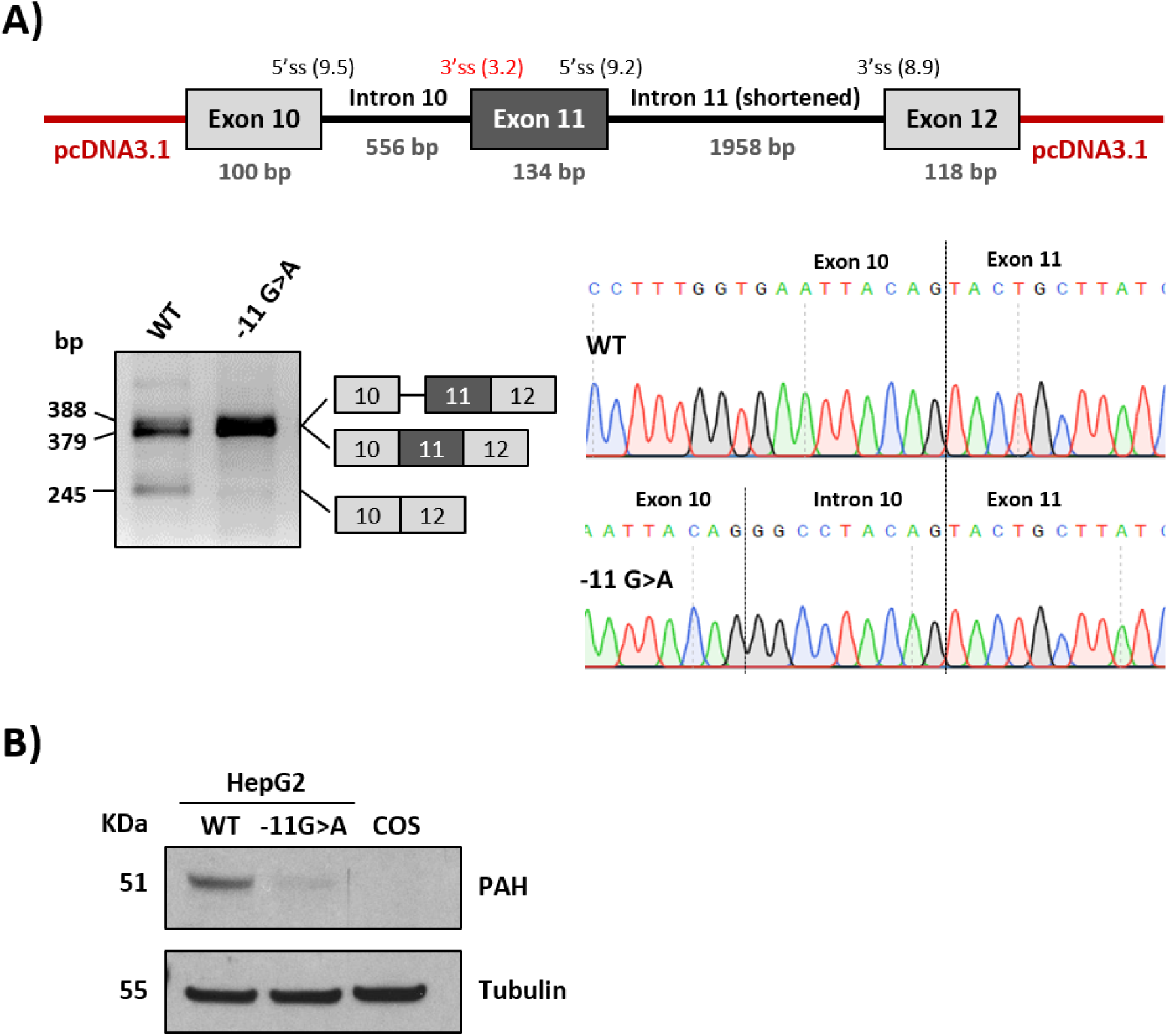
Functional analysis of the c.1066-11G>A *PAH* variant. **A)** Schematic representation of minigene construction in the pcDNA3.1 vector, RT-PCR after transfection in Hep3B cells with wild-type (WT) and mutant minigenes and Sanger sequencing. **B)** Western Blot for immunodetection of PAH from HepG2 WT and mutant cell lines. COS cells were used as negative control. Tubulin was used as loading control.

### Lack of PAH in c.1066-11A variant-engineered HepG2 cells

We next studied the expression of the mutant protein in a human hepatic context. To that end, we used CRISPR/Cas9 to introduce the c.1066-11G>A variant in HepG2 cells. After confirming the presence of the variant in homozygous fashion in the selected clone and the resulting splicing defect by RT-PCR (data not shown), western blot analysis showed almost no detectable immunoreactive protein (Figure 1B), which correlated with total absence of catalytic PAH activity (non-detectable).

### The humanized *Pah* c.1066-11A mouse model exhibits a classical PKU phenotype

The *KI Pah* c.1066-11A mouse model was generated using CRISPR/Cas9 and homology driven repair (HDR) gene editing. Examination of intron 10 sequence revealed no similarity between murine and human sequences, as expected. In fact, the GGGG sequence where the variant is located is not present in the murine sequence. Therefore, our strategy was to replace murine intron 10 sequence, adjacent to exon 11, with the human sequence carrying the variant. However, humanization of splicing sequences in a murine genomic context can have undesirable effects on the splicing outcome, as previously reported (15, 16). In order to confirm the correct recognition of the human sequence in the murine context and the recapitulation of the splicing defect, we first validated the strategy using minigenes, reproducing the construct designed for the mouse gene editing experiment. Murine exon 10, a partially humanized intron 10, murine exon 11, a shortened intron 11 and exon 12 were cloned in the pcDNA3.1 vector. This minigene in its WT version resulted in a correctly spliced exon 10-11-12 transcript and a transcript corresponding to exon 11 skipping. The mutant (c.1066-11A) hybrid minigene accurately recapitulated the splicing defect, with a sole transcript in which we confirmed the insertion of 9 extra nucleotides due to the use of the newly created splice site (Supplementary Figure 1A, B).

The humanized mouse model homozygous for the c.1066-11A variant was generated by CRISPR/Cas9 gene editing using the same design and the heterozygous founder mouse was identified by PCR genotyping (Supplementary Figure 1C) and subsequent sequencing analysis. The founder mouse was backcrossed to C57BL6/J background up to 10 generations. Mating of heterozygous mice yielded the expected mendelian ratios of wild-type (WT), heterozygous and homozygous *Pah* c.1066-11A offspring. In all experiments with the newly generated *Pah* c.1066-11A mice, age and sex-matched WT littermates were used as the control group. RT-PCR from liver RNA samples of homozygous mice confirmed the splicing defect resulting in the aberrant insertion of extra 9 nucleotides in the mature mRNA (Supplementary Figure 1D, E). Normal *Pah* mRNA levels were confirmed by RT-qPCR analysis (data not shown).

*Pah* c.1066-11A mice exhibited high mortality at weaning and lethal seizures at 6-7 months of age, thus resulting in reduced survival (Figure 2A). They display reduced body weight and the fur hypopigmentation characteristic of mouse PKU models (Figure 2 B, C). Besides, most *Pah* c.1066-11A mice exhibit microphthalmia (Figure 2C) which has previously been reported in other PKU models with a severe phenotype and correlates with the ocular abnormalities observed in untreated PKU patients (17, 18).

**Figure 2.**
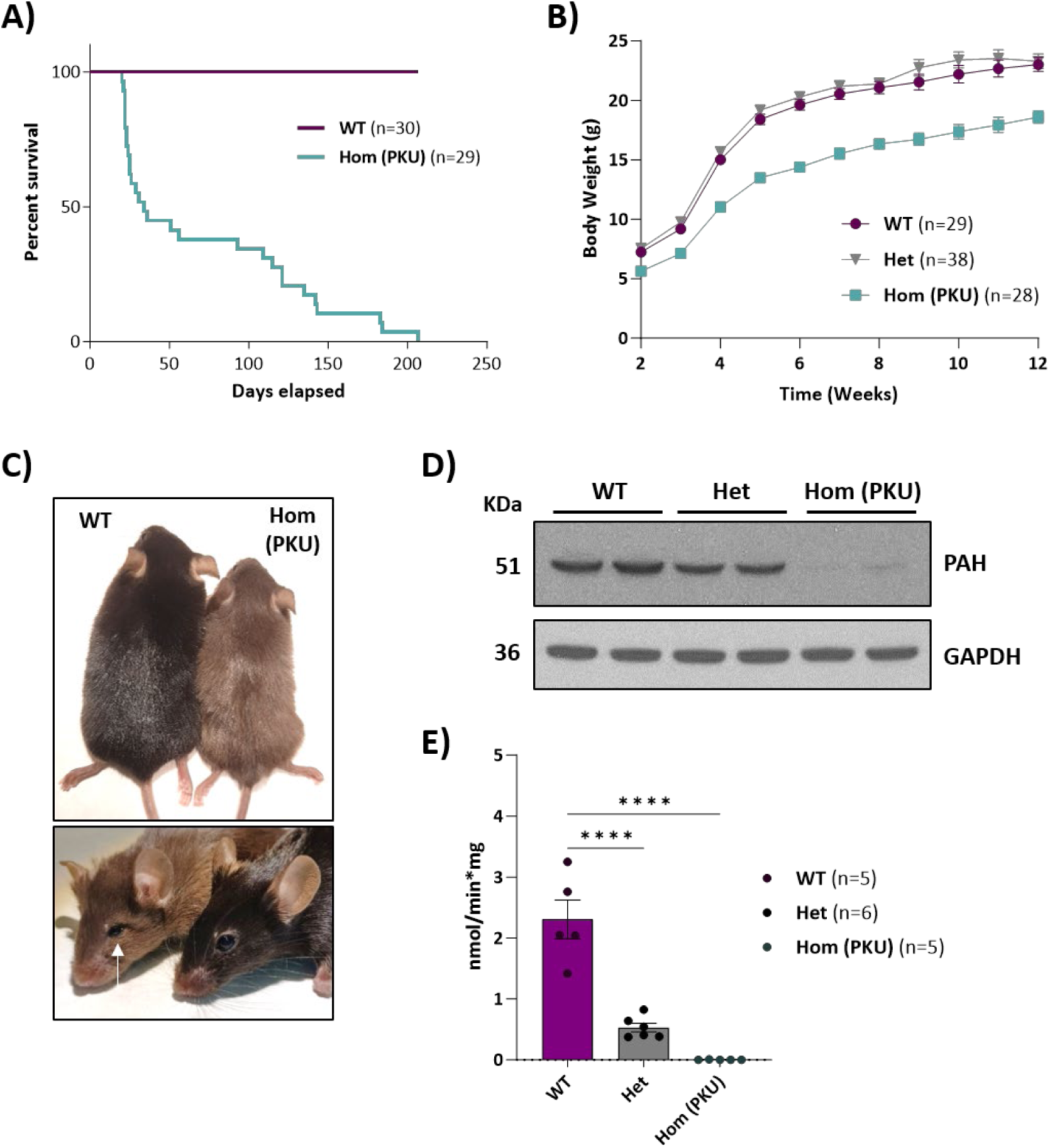
Characterization of *Pah* c.1066-11A (PKU) mice. **A)** PKU mice survival. **B)** Body weight distribution by genotype during 12 weeks. **C)** Hypopigmentation (12 weeks-old mice) and microphthalmia (20 weeks-old mice) in homozygous PKU mice compared with wild-type (WT). **D)** Western Blot for immunodetection of PAH from WT, heterozygous (Het) and PKU mice livers. GAPDH was used as loading control. **E)** Overview of PAH specific activity in liver lysates. Statistical analysis performed by one-way ANOVA (\*\*\*\**P*<0.0001). All plots show mean values ± SEM.

Western blot analysis in liver samples showed nearly undetectable hepatic PAH protein in *Pah* c.1066-11A PKU mice (Figure 2D). Accordingly, lack of hepatic PAH activity was confirmed (Figure 2E). Heterozygous mice exhibited slightly reduced levels of PAH protein (76%) and intermediate-low PAH activity (23%).

Biochemical analyses revealed high blood serum Phe (1885.7±71.2 μM *versus* 59.3±2.5 μM for WT) and low Tyr levels (28.8±1.6 μM *versus* 68.2±4.4 μM for WT), corresponding to severe, classical PKU. Female mice exhibited higher blood Phe levels as compared to males (2126.8±35 *versus* 1135 ± 53 µM). Sex differences in blood Phe levels have been previously reported in PKU mouse models (19), although they have not been described in PKU patients. In brain, Phe levels were also high in both sexes of the *Pah* c.1066-11A PKU model (153.4±3.9 μM *versus* 13.6±0.7 μM for WT) and Tyr levels were low (6.4±0.3 μM *versus* 14.6±1.2 μM for WT). Among all the remaining analysed amino acids, cystine was found significantly elevated in the *Pah* c.1066-11A mice serum, and lower levels were found for tryptophan, asparagine, glutamine, glycine, proline, alanine, methionine, ornithine, lysine and arginine (Supplementary Table 1). In brain, serine, glycine, lysine, histidine, and cystathionine were found elevated while glutamine, leucine and isoleucine were found in reduced levels in *Pah* c.1066-11A mice (Supplementary Table 2). No significant differences were observed between males and females.

Monoamine neurotransmitters and related metabolites (dopamine, serotonin, DOPAC, HVA and 5-HIAA) were significantly lower in brain of homozygous *Pah* c.1066-11A mice of both sexes as compared to WT mice (Figure 3).

**Figure 3.**
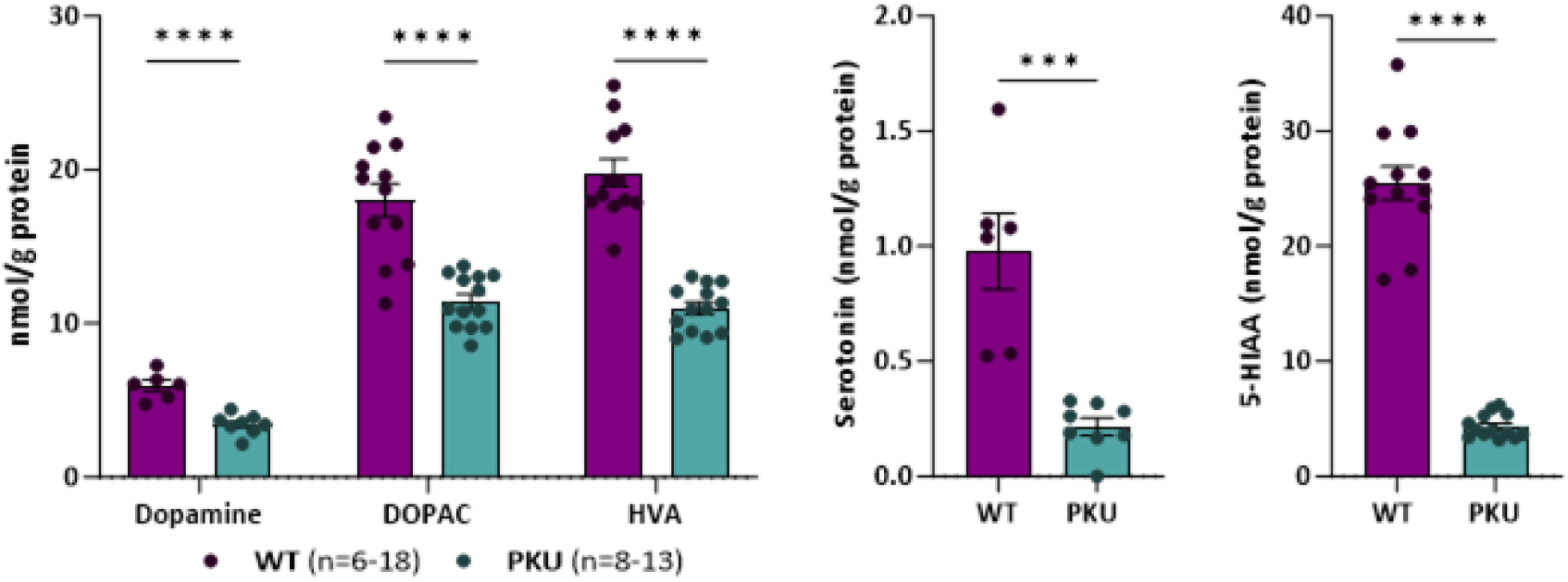
Brain neurotransmitters and related metabolites levels in *Pah* c.1066-11A (PKU) mice. Measures of neurotransmitters dopamine, 3,4-Dihydroxyphenylacetic acid (DOPAC), homovanillic acid (HVA), serotonin and 5-hydroxyindole acetic acid (5-HIAA) in brain lysates by HPLC. Statistical analysis performed by unpaired t test (\*\*\**P*<0.001, \*\*\*\**P*<0.0001). Data are presented as mean ± SEM.

We also examined the response to BH_4_ treatment in the novel PKU mouse model. BH_4_ was administered at 10 mg/kg body weight twice a day for four consecutive days and Phe levels measured in dried blood spots sampled daily after 3 hours fasting. As seen in Supplementary Figure 2, no significant changes in blood Phe content were detected in BH_4_ treated PKU mice compared to those receiving placebo (the same solution without BH_4_).

### Humanized *Pah* c.1066-11A mice exhibit neurological alterations

Different behavioural tests were carried out in 12–14-week-old mice (Figure 4). First, mice were analysed in the open field test to evaluate their general locomotor activity. *Pah* c.1066-11A mice showed decreased total ambulatory distance covered, as well as decreased average velocity, indicative of a general hypoactivity phenotype (Figure 4A). Curiously, the decreased ambulatory activity is more pronounced in the centre of the arena, what might be suggestive of anxiety-like behaviour (Figure 4A). Motor coordination and neuromuscular fitness were also assessed in the rotarod and inverted grid tests, respectively. In the rotarod, *Pah* c.1066-11A mice showed a significant decrease in their latency to fall from the accelerating rod, compared to WT mice, thus evidencing a deficit in motor coordination (Figure 4B). The latency to fall was also markedly decreased in the inverted grid test (Figure 4C). Interestingly, *Pah* c.1066-11A mice also showed hindlimb clasping (Figure 4D), a common phenotype in several mouse models of neurodegeneration, including certain cerebellar ataxias and striatal disorders. Furthermore, *Pah* c.1066-11A mice were unable to perform the vertical pole test; they immediately fell down, as they were not able to hold on to it, thus further revealing a deficit in motor coordination and neuromuscular fitness. Finally, *Pah* c.1066-11A mice were also subjected to the social approach test. In general, *Pah* c.1066-11A mice behaved similar to WT mice in this test. However, when exploring the occupied cage, *Pah* c.1066-11A mice showed a decreased number of physical contacts with the other mouse, what might be indicative of diminished sociability (Figure 4E).

**Figure 4.**
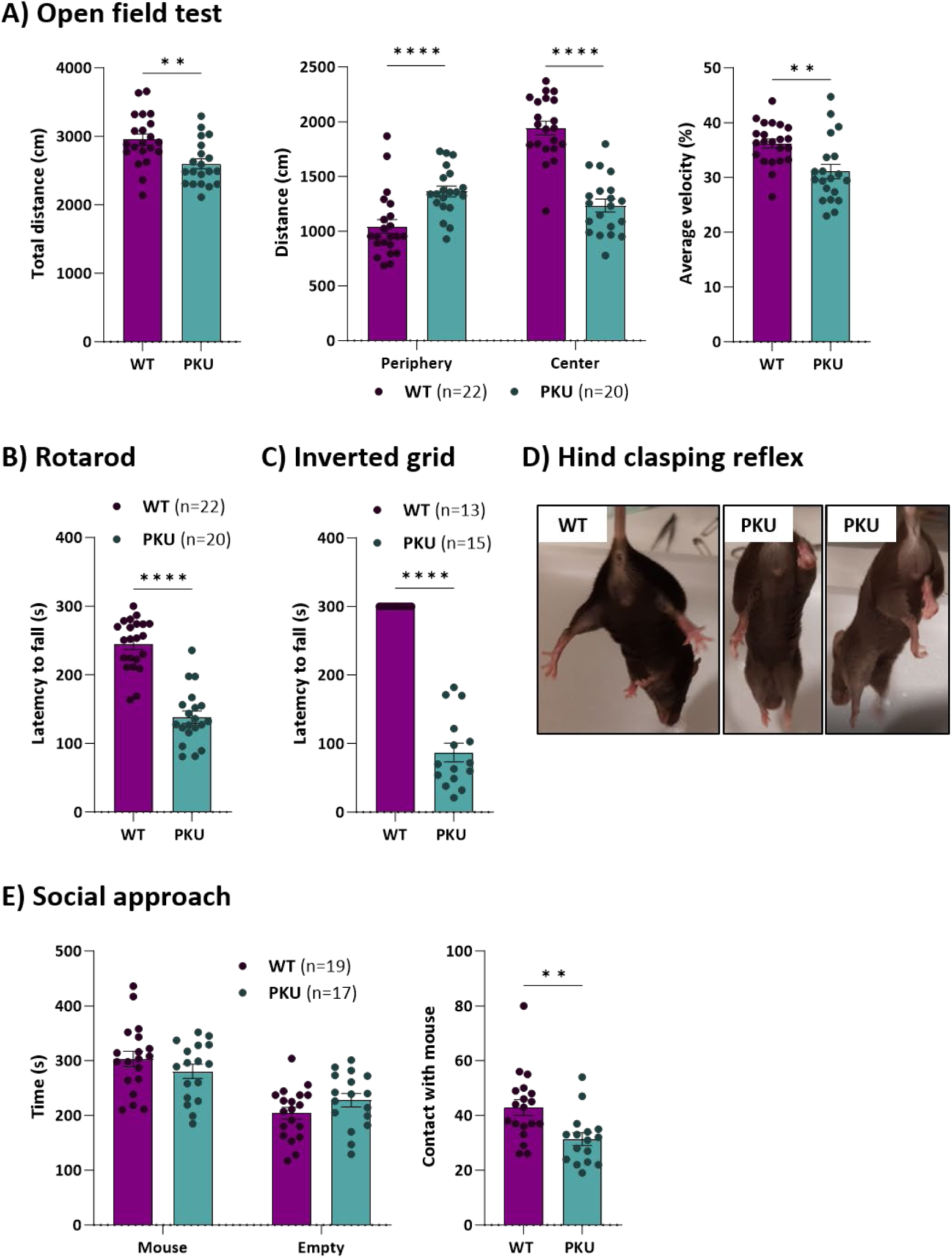
Behavioural performance of *Pah* c.1066-11A (PKU) mice. **A)** Distance travelled and velocity in open field test. **B)** Hind-limb clasping. **C)** Latency to fall off the rotarod. **D)** Latency to fall off the inverted grid test. **E)** Time spent interacting with empty cage and unfamiliar mouse in social approach test. Statistical analysis performed by unpaired t test (\*\**P*<0.01, \*\*\*\**P*<0.0001). Data are presented as mean ± SEM.

We next examined selected neuronal and glial markers by immunohistochemistry analysis in different brain regions of 20 weeks-old *Pah* c.1066-11A mice, as compared to age matched WT mice. No differences were found when staining for the neuronal marker NeuN (data not shown). On the contrary, both GFAP and Iba1 staining signals were significantly increased in *Pah* c.1066-11A mice, revealing changes in the morphology of both astrocytes and microglial cells, as shown for the dentate gyrus of the hippocampal formation (Figure 5). In contrast, Olig2 staining was significantly reduced indicating a deficit in oligodendrocytes, as shown for cortex, striatum and corpus callosum (Figure 6A), which is in line with the reported myelinization deficits in PKU (20). Indeed, we could confirm decreased myelinization in brains of the *Pah* c.1066-11A mice by electron microscopy, with decreased thickness of the myelin sheaths and loss of myelinated axons in the lateral tract of the spinal cord (Figure 6B).

**Figure 5.**
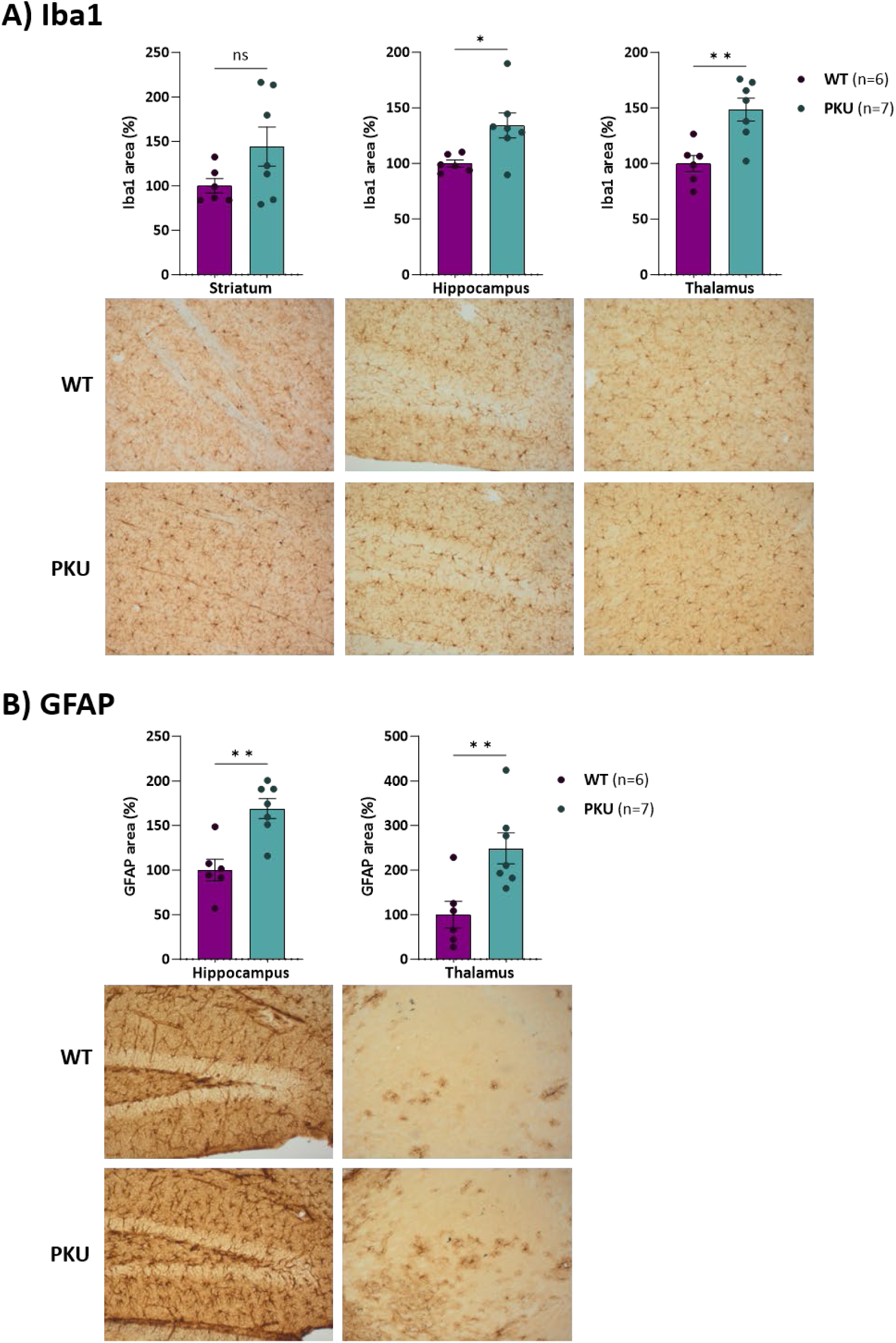
Analysis of different types of neuronal markers. Immunohistochemistry brain image of Iba1 **(A)** and GFAP **(B)** in WT and PKU mice. Statistical analysis performed by unpaired t test (\**P*<0.05, \*\**P*<0.01, ns, not significant). Data are presented as mean ± SEM.

**Figure 6.**
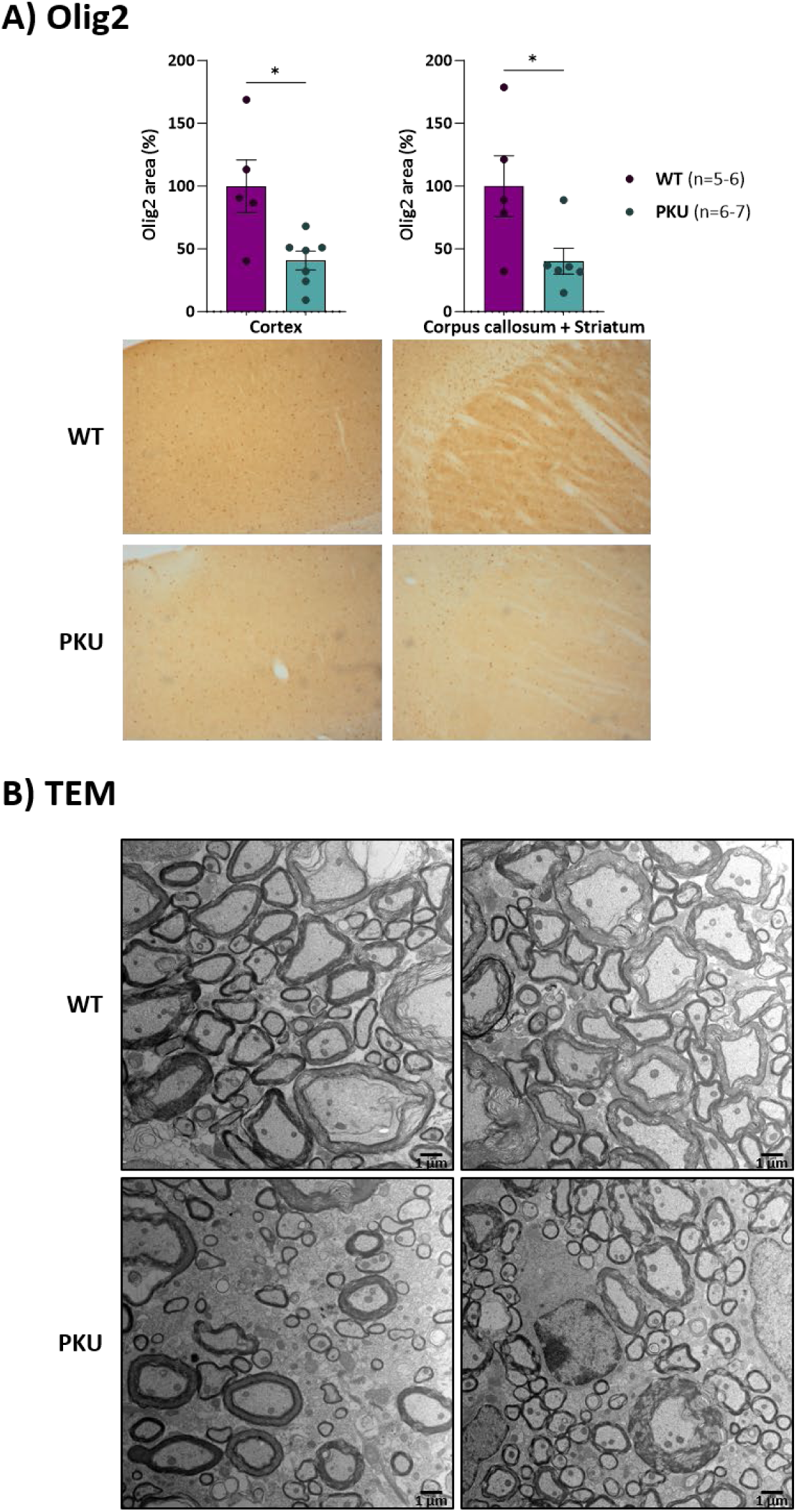
Myelinization defects. **A)** Immunohistochemistry brain images of Olig2 in Wt and PKU mice. Statistical analysis performed by unpaired t test (\**P*<0.05). Data are presented as mean ± SEM. **B)** Transmission electron microscopy (TEM) images of lumbar transverse sections of the spinal cord of two different WT and *Pah* c.1066-11A (PKU) mice.

### The pQ355_Y356insGLQ mutant PAH protein is highly unstable

Structural predictions for the pQ355_Y356insGLQ mutant PAH protein resulting from the c.1066-11G>A variant indicate loss of H-bonds relevant for the correct position of α-helix 11 and steric clashing of the inserted Leu with L333 (Supplementary Figure 3). AlphaFold predicts an additional turn of the helix in the mutant protein but introduces a torsion in order to maintain the interactions of Y356 as in the WT. Therefore, it is probable that the insertion of 3 amino acids hampers the correct arrangement of residues in this region, thereby affecting protein folding. This correlates with the low levels of detected protein *in vitro* and *in vivo*.

Mutant forms of the PAH protein undergoing misfolding and/or aggregation have been reported to be targets for ubiquitin-mediated proteasomal degradation or selective autophagy (21). Accordingly, we examined the impact of the expression of the mutant p.Q355_Y356insGLQ protein in those processes, as well as on the levels of the co-chaperone DNAJC12 and of HSP70 that participate in the correct folding and stabilisation of the PAH protein (22, 23). Western blot analysis in mouse liver lysates showed no significant changes in the ubiquitination profile between WT and *Pah* c.1066-11A mice samples. However, both DNAJC12 and HSP70 protein levels were decreased (Figure 7A). No significant differences in gene expression levels were detected for the corresponding *Dnajc12* and *Hsp70* genes by RT-qPCR analysis (data not shown). Overall, the results could be indicative of some degree of intracellular co-aggregation of the molecular chaperones with the mutant protein.

**Figure 7.**
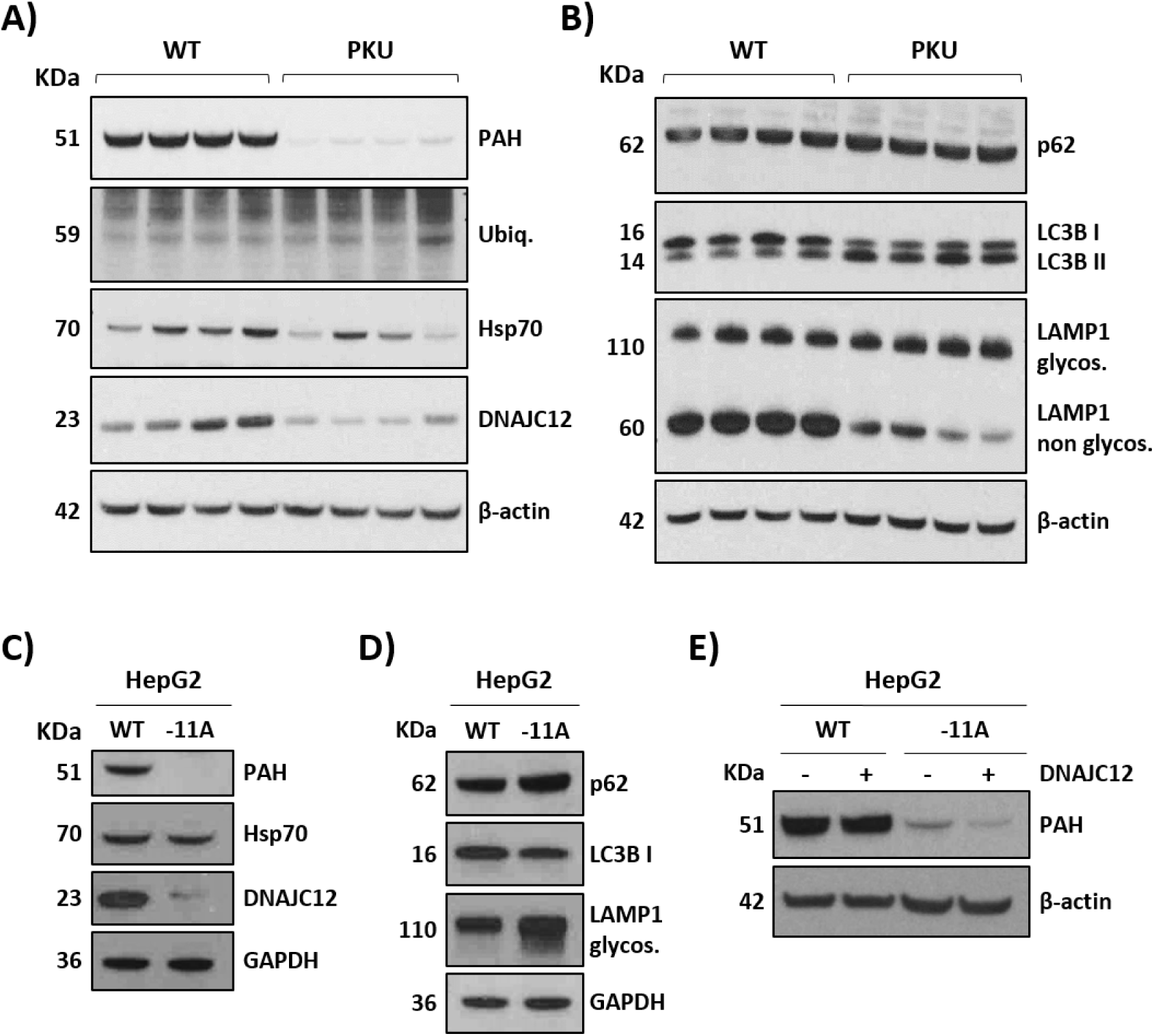
Evaluation of the effect of the c.1066-11G>A variant on PAH folding and stability. **(A, B)** Western blot analysis of PAH, ubiquitin and DNAJC12 and HSP70 co-chaperones and autophagy markers LAMP1, LC3BII and p62 in liver samples from 12-week-old mice. **(C, D)** Western blot analysis in HepG2 cells. **E)** PAH levels detected by western blot in HepG2 cells 72 h after transfection with 2 µg of the plasmid coding for DNAJC12. GAPDH or β-actin were used as loading controls.

PAH protein aggregates may be processed by selective autophagy rather than by the ubiquitin-dependent proteasome system (7, 21). We detected altered levels of autophagic markers in *Pah* c.1066-11A mice livers, with higher levels of glycosylated LAMP1 protein and an increase in the conversion of LC3-I to the lower migrating form LC3-II (Figure 7B).

To confirm these results in a human genetic context, we analyzed these markers in the gene edited HepG2 cells carrying the c.1066-11A variant. DNAJC12 protein levels were also decreased, and we could detect decreased levels of LC3BI/II and an increase in glycosylated LAMP1 (Figure 7C, D).

We recently showed that coexpression of DNAJC12 with different PAH mutant constructs in COS7 cells results in a mutation-dependent effect, with an increase in protein levels and activity for some unstable mutants (23). To examine the effect of DNAJC12 overexpression on the mutant p.Q355_Y356insGLQ protein, we transfected the edited HepG2 cells with the pReceiver/DNAJC12 plasmid (23) and analysed PAH protein levels after 72h. The results showed no significant change in the low levels of mutant PAH protein present in the edited cells (Figure 7E).

## Discussion

The development of personalized medicine for genetic diseases requires preclinical testing in appropriate models. *PAH* variant c.1066-11G>A, a splicing defect resulting in an in-frame insertion of 3 amino acids, is the second most common pathogenic variant causing PKU worldwide, prevalent in Southern Europe and Latin America. Innovative RNA-based therapeutics such as splice switching antisense oligonucleotides (24) or HULC lncRNA mimics (8), gene editing approaches (9, 10, 25, 26) or use of specific pharmacological chaperones (27, 28), are theoretically applicable to target either the mutant pre-mRNA or the resulting protein, with the aim of correcting the pathological phenotype of this frequent variant. Due to the limitation in access to patients’ tissues where the *PAH* gene is expressed (liver and to a lesser extent, kidney) the splicing defect was studied using minigenes and in a gene edited HepG2 cell line homozygous for the c.1066-11G>A variant, in which almost no detectable PAH protein and activity were detected. While *in vitro* studies can be performed in cellular models, for a successful clinical translation of potential therapies, a specific *in vivo* model is needed, in which the gene is expressed in its natural context. This allows to evaluate efficacy in target gene engagement, functional protein expression, downstream functional readouts, therapeutic compounds’ uptake and biodistribution, and to perform potential toxicology studies.

As mouse and human intronic sequences are non-homologous, in the present work we have employed CRISPR/Cas9 technology to replace the murine sequence of *Pah* intron 10, proximal to exon 11, with the human sequence carrying the c.1066-11G>A variant. We verified that no modifications of the predicted off-target sites were detected in the humanized mouse and that the genome-editing approach did not interfere with *Pah* gene expression in liver, where precise recapitulation of the splicing defect was confirmed. As expected, no functional PAH protein was detectable in liver resulting in a phenotype resembling the classical form of PKU, with Phe levels above 1200 μmol/L and corresponding low Tyr and Trp levels. Elevated Phe and low Tyr levels were also detected in brain samples of homozygous *Pah* c.1066-11A mice, along with significant dopamine and serotonin deficiencies, which have been implicated in the pathogenesis of the neurological, behavioural and cognitive deficits observed in untreated patients and in PKU mouse models (19, 29). Correspondingly, *Pah* c.1066-11A mice exhibit locomotor and behavioural abnormalities, as well as altered hind-limb clasping which is a common feature of different neurological and neurodegenerative diseases (30). This is associated with impaired brain connectivity, possibly mediated by changes in monoaminergic transmission to different brain regions.

Neuropathological evaluation of different PKU mouse models have shown white matter alterations (20, 31), decreased synaptic connections but normal neuron numbers (32), in agreement with brain histology reports from untreated PKU patients (33, 34). Classical PKU patients, left untreated, exhibit intellectual disability, developmental delay, epilepsy, autistic features, and movement disorders. Our results obtained with the novel PKU mouse model created are in agreement with these observations. Immunohistochemistry analysis in the *Pah* c.1066-11A mice confirmed an altered brain physiology, with changes in the morphology of astrocytes and microglia, decreased levels of oligodendrocytes and normal neuronal content. Hypomyelination is probably the most important factor in the observed motor impairment with neurological alterations in PKU (34). Elevated Phe levels inhibit 3-hydroxy-3-methylglutaryl coenzyme A reductase (HMGR), thus reducing cholesterol biosynthesis in the PKU brain (35). Cortex and corpus callosum appear to be specifically vulnerable to high Phe levels (34), consistent with our Olig2 immunohistochemistry observations. Other factors possibly contributing to the observed hypomyelination in PKU have been put forward: epigenetic modifications, oxidative stress, defects in calcium homeostasis or altered protein synthesis at the myelin sheath (31).

Our results demonstrate that *Pah* c.1066-11A mice recapitulate PKU disease biology, with biochemical and clinical alterations observed in the most severe classical PKU phenotype, consistent with almost no detectable PAH protein and activity in liver. It thus represents a useful avatar mouse model for a frequent PKU variant, for physiopathology studies and for assessing efficacy of various innovative therapeutic modalities. In this sense, for splicing variants, our group has efficiently used splice-switching AONs to correct the resulting splice defects (36–38). The c.1066-11G>A variant is in close proximity to the natural 3’ splice acceptor site, hindering the design of an AON that could effectively block the novel splice site created by the variant, without causing additional mis-splicing defects such as exon skipping. Indeed, the insertion of 9 extra nucleotides due to the similar variant c.3191-11T>A in the *ABCA4* gene causing Stargardt disease, could not be corrected by the single AON that could be designed, taking into account the limitations imposed by the target sequence region (39). Other strategies targeting binding sites of splice factors involved in the negative regulation of exon 11 splicing can be envisaged. Future studies will test the feasibility of AON-mediated splice correction for the c.1066-11G>A variant.

Alternatively, gene editing approaches, which have shown to be effective in correcting PKU variants in different mouse models (9, 10, 25, 26), could provide a novel therapeutic approach for PKU patients with this specific variant. In this sense, the created avatar PKU mouse model with the humanized mutant (c.1066-11A) intron 10 fragment of the endogenous *Pah* gene is a suitable preclinical model for assessing the efficacy of adenine base editing as a therapeutic intervention, which could be applicable to a high number of PKU patients worldwide.

In this study we confirmed that the mutant p.Q355_Y356insGLQ PAH protein resulting from the c.1066-11G>A variant was found to be highly unstable, in both *in vitro* (HepG2) and *in vivo* (mouse) models generated in this work. Previous studies using a prokaryotic expression system showed that the mutant protein fused with maltose binding protein was recovered exclusively as high molecular weight aggregates (12). In this work, the reduction of immunodetected levels of DNAJC12 and, to a lesser extent, of HSP70, indicate a possible coaggregation of the mutant PAH protein with these chaperones. This has been described for several unstable human *PAH* variants and for the highly unstable *Pah* variant p.V106A present in the Enu1/1 mouse model (21). Our results support a role of DNAJC12 in the processing of misfolded aggregation-prone p.Q355_Y356insGLQ mutant protein by the autophagy system. Overexpression of DNAJC12 in the edited HepG2 cells did not counteract the stability defect. Our group recently described a mutation-dependent effect of the overexpression of DNAJC12 on missense PAH mutant proteins transiently expressed in COS7 cells; for some there was an increase in protein and activity levels while for others the effects on stability were aggravated (23). The outcome may depend on the exact pathogenic mechanism of each variant, with DNAJC12 guiding the Hsp70 machinery towards the specific folding or degradation of the client protein. Further studies are necessary to reveal the interactions between the mutant PAH protein and the quality control system and the exact mechanism implicated in its degradation.

In summary, our work has provided a translationally relevant murine model of classical PKU resulting from a frequent PAH variant, as well as the corresponding human hepatoma cellular model, to be used as useful experimental tools to advance research in PKU pathophysiology and personalised therapies.

## Methods

### Cell culture and treatments

Human hepatoma cell lines were grown in Minimum Essential Medium (MEM, Sigma Aldrich) supplemented with 5% fetal bovine serum (FBS), 1% glutamine and 0.1% antibiotic mix (penicillin/streptomycin) under standard cell culture conditions (37°C, 95% relative humidity, 5% CO2). They were routinely tested for mycoplasma contamination.

### Minigenes construction

For evaluation of *in vitro* splicing, a minigene construction including *PAH* exon 10, full intron 10, exon 11, 1958 bp of intron 11, and exon 12 cloned in pcDNA3.1+ (40) was used. A mutant minigene with the c.1066-11G>A variant was generated by site-directed mutagenesis with QuikChange Lightning Kit (Agilent Technologies, Santa Clara, CA, USA). Sanger sequencing confirmed the identity of the constructs. To study the effect of humanizing 90 bp of the 3’ end of intron 10 of the mouse *Pah* gene containing the c.1066-11G>A mutation for the subsequent generation of the murine model, a hybrid minigene was constructed with this design and including from *Pah* exon 10 to exon 12, with a shortened intron 11. The sequence of interest, with a total length of 2532 base pairs, was synthesized and cloned into pcDNA3.1+ between the restriction enzymes KpnI and XhoI, using the GeneArt service (ThermoFisher Scientific, USA).

### Transient transfection and splicing analysis

For minigene assays, Hep3B cells were seeded in six-well plates at a density of 4x10^5^ in 2 ml 10% MEM and grown overnight. Cells were transfected with a total DNA amount of 2 μg per well using lipofectamine 2000 (Invitrogen) and harvested by trypsinization after 48 h. Total RNA was extracted using Trizol Reagent (ThermoFisher) and phenol-chloroform. cDNA synthesis was performed using NZY First-Strand cDNA Synthesis Kit (NZYtech). Splicing analysis was carried out by PCR amplification with FastStart Taq Polymerase (Roche) using specific primers (Supplementary Table 3). End-point PCR amplification products were analyzed by 3% agarose gel electrophoresis.

### Generation of the *KI* cell line HepG2 *PAH* c.1066-11A

HepG2 cells were obtained from ATCC company (HB-8085). The *KI* cell model was generated via CRISPR/Cas9 using Breaking Cas software (https://bioinfogp.cnb.csic.es/tools/breakingcas/) (41) to design guides (crRNA) with high specificity to introduce the c.1066-11G>A variant in the PAH gene and to identify potential off-targets (Supplementary Table 4). To generate the sgRNA complexes, crRNA and tracrRNA (Integrated DNA Technologies, IDT) were mixed in equimolar concentrations. The ribonucleoprotein complex (RNP) was formed by the combination of 24 pmoles of Cas9 (Streptococcus pyogenes, IDT) and 24 pmoles of sgRNA. Approximately 6x10^5^ HepG2 cells per well were seeded in a 6-well plate with the RNP complex and 2.4 pmoles of the DNA template for homology driven repair (HDR) (Ultramer oligonucleotide, IDT) (Supplementary Table 4) using Lipofectamine CRISPRMAX™ (ThermoFisher Scientific Waltham). After 48h, the culture media was changed to fresh media. Three days post-transfection, cells were dispersed at low density into 150 mm dishes in MEM medium. About 15-20 days later, large colonies were picked and expanded. Genomic DNA extraction of different clones was carried out using QIAamp DNA Mini Kit (Qiagen). DNA concentration was measured using Nanodrop One spectrophotometer (ThermoFisher Scientific). The region surrounding the desired change was amplified by PCR and subsequently digested with the restriction enzyme DdeI (New Englands Biolabs) to detect positive clones, which were confirmed by Sanger sequencing. The three predicted highest risk off-target sites were also amplified using specific primers (Supplementary Table 4) and sequenced.

### Generation of humanized *KI* mouse *Pah* c.1066-11A

PKU humanized mice were generated in a 50% C57BL/6J and 50% CBA/CA background via CRISPR/Cas9-mediated gene editing, by the institutional core facility (CNB-CBM mouse transgenesis). The CRISPR sgRNA (300 ng/mL) was co-injected into fertilized oocytes with HiFi Cas9 protein (600 ng/mL) and DNA Template (400 ng/mL) (Supplementary Table 5), all of them synthesized and purchased from IDT. The DNA template harbored 90 base pairs of the 3’ end of human *PAH* intron 10 containing the c.1066-11G>A pathogenic variant flanked by mouse *Pah* intron 10 sequence. One animal was found to be heterozygous for the c.1066-11G>A variant and the correct wild-type sequence from a total of 14 pups born. Confirmation of the on-target and the verification of the absence of the four highest risk off-target sites were performed from genomic DNA tail biopsies by PCR amplification with specific primers (Supplementary Table 5) and subsequent Sanger analysis.

### Mouse husbandry and colony expansion

Animal housing and maintenance protocols followed the local authority guidelines. Mice were maintained on standard chow. All the experiments were carried out in a pathogen-free environment at the animal facility of the Centro de Biología Molecular Severo Ochoa, in accordance with the Spanish Law of Animal Protection. Food and water were available ad libitum, and mice were maintained in a temperature-controlled environment on a 12/12 h light-dark cycle.

The colony was backcrossed up to F10 with wild-type mice (C57BL/6J). Heterozygous mice were used for breeding and the resultant homozygous mutant *Pah* c.1066-11A and wild-type (WT) mice have been used in the experiments presented in this work. All mice used were adults (3–5 months-old) and males and females were included in all experiments. Body weight was monitored every week. At the time/age stated for each experiment, mice were sacrificed in a carbon dioxide chamber and perfused with 50 ml of NaCl. Immediately after, brain and liver were surgically excised, snap-frozen in liquid nitrogen and stored at −80 °C.

### BH_4_-loading test

A treatment solution of BH_4_ (Sigma Aldrich, T4425) (2mg/ml BH_4_, 2% ascorbic acid, and 10% DMSO; in a dosage of 20 mg BH4/kg body weight) or placebo solution (2% ascorbic acid and 10% DMSO), was injected daily intraperitoneally into PKU mice (n = 5 mice per group) in 2 doses for 4 days. Blood samples were collected onto filter paper cards after 3 h fasting, before the first injection and during the 4 days, and Phe was quantified by fluorimetry using Neonatal Phenylalanine kit (Labsystems Diagnostics).

### Amino acid and neurotransmitters determination in brain

For analysis of amino acid concentrations in brain, half-brains were homogenized using Ultra-Turrax device (IKA) in 5 volumes/weight 10% trichloroacetic acid containing norvaline (150 µM). The homogenates were clarified by centrifugation at 500 g, 5 min at 4⁰C and supernatants stored at −80°C until analysis by ion exchange chromatography with ninhydrin. For neurotransmitters analysis half-brains were processed by adding 3 volumes/weight of a lysis buffer solution (50 mM Tris-HCl pH 7.5, 150 mM NaCl, 0.1% Triton X-100, 1 mM EDTA, and 10% glycerol) with protease inhibitor. Analysis of biogenic amine neurotransmitters metabolites, 5-hydroxyindolacetic acid (5-HIAA), homovanillic acid (HVA), 3-metoxy-hydroxy-phenylglycol (MHGP), 3-O-methyldopa (3-OMD), 5-hydroxytryptophan (5-OH-TRP), dihydroxyphenylacetic acid (DOPAC), dopamine and serotonin in brain lysates was performed by ion pair high-performance liquid chromatography (HPLC) with electrochemical detection (ESA, Coulocem III) as reported (42). Briefly, samples were deproteinized with VWR Centrifugal Filters, diluted 1:2 in the chromatography mobile phase and 50µl were injected into HPLC. Measured brain homogenate monoamine neurotransmitter concentrations were corrected for the protein content of the homogenate and expressed as pmol/mg protein.

### Amino acid determination in serum

In order to obtain serum specimens, whole blood was collected by cardiac puncture and transferred to an Eppendorf tube. Subsequently, blood samples were left for 1 h at 37°C and at 4°C overnight to facilitate coagulation, followed by two consecutive centrifugation steps (1200 g, 5 min, and 4 °C) after which the respective supernatants were pipetted out to a clean tube. The isolated serum fractions were stored at −80 °C and amino acids measured by ion exchange chromatography (visualized with ninhydrin).

### PAH enzymatic activity assay

For PAH enzyme activity determination, cells were lysed by adding Na-Hepes buffer (50 mM HEPES pH 7.0, 500 mM NaCl) followed by three freeze-thawing cycles. Proteins were collected in the supernatant fraction after centrifugation 10 min at 16,000 g at 4°C. Liver lysates were prepared with Na-Hepes buffer, shredding the tissue with ultra-turrax and centrifuged at 16000 g for 30 min. PAH activity in the homogenates was measured at room temperature (RT≈25°C) using 10 μg of total protein in each assay, with 1 mM L-Phe in Na–Hepes pH 7.0, containing catalase (0.1 mg/ml). After 5 min preincubation at 25°C, ferrous ammonium sulfate (100 μM) was added, and the reaction was triggered after 1 min by adding 75 μM BH_4_ and 5 mM DTT (final concentrations in the assay). The reactions were allowed to run for 30 min at RT and stopped with 3 volumes of 12% perchloric acid, followed by incubation at 4°C for 1 h and at -20°C overnight. The mix was centrifuged at 16000 g, 15 min at 4°C and supernatants stored at −20°C until analysis. Under these conditions, PAH activity was linear to the amount of protein in the extracts. L-Tyr formed was quantified by HPLC with fluorometric detection as described previously (43).

### Cells, liver and brain protein analysis

Cells were harvested by trypsinization and treated with RIPA lysis buffer (10 mM Tris-HCl 7.5, 150 mM NaCl, 0.1% Triton, and 10% glycerol), protease and phosphatase inhibitors. Samples were subjected to three freeze-thawing cycles and proteins were collected in the supernatant fraction after centrifugation at 16,000 g for 10 min at 4°C. Mouse liver and brain proteins were isolated by disrupting the tissue in lysis buffer (50 mM Tris-HCl pH 7.5, 150 mM NaCl, 0.1% Triton X-100, 1 mM EDTA, and 10% glycerol) using ultra-turrax followed by centrifugation at 16,000 *g* for 30 min at 4°C. Protein concentration in the supernatant was measured using the Bradford method (Bio-Rad Laboratories).For western blot analysis, equal amounts of protein (50 µg) were loaded into 4-12% SDS-polyacrylamide gels. Proteins were transferred into a nitrocellulose membrane using the iBlot Dry Blotting System (Thermo Fisher Scientific). For the analysis of total proteins, membranes were blocked for 1 hour with 5% nonfat milk, in 0.1% TBS-tween and incubated overnight with the corresponding primary antibody: PAH (1:1000, Santa Cruz Biotechnology, sc-271258), Ubiquitin (1:1000, Santa Cruz Biotechnology, sc-8017), Hsp70 (1:1000, Novus Biological, NB110-61582), DNAJC12 (1/1000, Abcam, ab167425), LC3B (1:1000, Cell Signaling Technology, #2775), LAMP1 (1:1000, Cell Signaling Technology, #3243) and p62/SQSTM1 (1:5000, Novus biologicals, H00008878-M01). Secondary antibodies were horse anti-mouse (1:2000, Cell Signaling Technology, 7076S) and goat anti rabbit (1:5000, Cell Signaling Technology,7074S). Antibody against GAPDH (1:5000, Abcam, ab8245) and β-actin (1:2000, Origene, TA811000S) was used as loading control. Enhanced chemiluminescence reagent (ECL, Cytiva) or SuperSignal West Femto (ThermoFisher) were used for protein detection. Band intensity for each protein was quantified with BioRad GS-900 Densitometer (Bio-Rad) and ImageLab program.

### mRNA quantification

For gene expression analysis, cDNA was obtained by retrotranscription of 500 ng of total RNA from mouse liver or HepG2 cells using NZY First-Strand cDNA synthesis kit (NZYTech). *Pah, Hsp70, Dnajc12*, genes were amplified with specific primers (Supplementary Table 1), using Perfecta SYBR Green FastMix kit (Quanta Biosciences) in a CFX Opus 384 (Bio-Rad). *Gapdh* was used as endogenous control and quantification was done using the 2^−ΔΔCt^ method.

### Immunohistochemistry

Brain hemispheres were extracted and fixed overnight in 4% paraformaldehyde, cryopreserved for 72h in 30% sucrose in PBS and included in optimum cutting temperature (OCT) compound (TissueTek, Sakura Finetek Europe, 4583), immediately frozen and stored at −80 °C until use. Using a cryostat (Thermo-Fisher Scientific) 30 µm sagittal sections were cut, collected and stored free floating at −20 °C in glycol containing buffer (30% glycerol, 30% ethylene glycol in 0.02 M phosphate buffer). Mouse sections were washed in PBS, immersed in 0.3% H_2_O_2_ in PBS for 45 min and blocked for 1h in blocking solution (PBS containing 0.5% fetal bovine serum, 0.3% Triton X-100 and 1% BSA) and then incubated overnight at 4°C with the primary antibody diluted in blocking solution. After washing, sections were incubated with biotinylated anti-mouse or anti-rabbit secondary antibody and then with avidin-biotin complex using the Elite Vectastain kit (Vector Laboratories, PK-6101 and PK-6102). Chromogen reactions were performed using diaminobenzidine (SIGMAFAST DAB, Sigma-Aldrich, D4293) for 10 min. Sections were mounted on glass slides and coverslipped with Mowiol (Calbiochem, 475904). An Axioskop2 plus (Zeiss) microscope with an DMC6200 camera (Leica) was used to capture the images.

### Transmission electron microscopy

Mice were completely anaesthetized with an intraperitoneal tiopental injection (100 µl/10 g), perfused transcardially with 50 ml of NaCl and then with a solution of 4% formaldehyde and 2 % glutaraldehyde in 0.1M phosphate buffer pH 7.4 Spinal cords were postfixed in the same solution for 2 h at room temperature and overnight at 4°C. Spinal cord lumbar fragments were postfixed in 1% osmium tetroxide in 0.1 M cacodylate buffer, dehydrated in ethanol and embedded in Epon-Araldite. Serial ultrathin sections of the lumbar transverse section of the spinal cord were collected on pioloform-coated, single-hole grids, and stained with uranyl acetate and lead citrate. The sections were analysed with a JEM-1010 transmission electron microscope (Jeol, Japan) equipped with a side-mounted CCD camera Mega View III from Olympus Soft Imaging System GmBH (Muenster, Germany). Myelinated fibers were sampled randomly in the white matter tracts of the spinal cord at a magnification of 3000X.

### Behavioral testing

#### Open field test

Locomotor activity was measured in clear Plexiglas® boxes measuring 27.5cm x 27.5cm, outfitted with photo-beam detectors for monitoring horizontal and vertical activity. Activity was recorded with a MED Associates’ Activity Monitor (MED Associates) and were analyzed with the MED Associates’ Activity Monitor Data Analysis v.5.93.773 software. Mice were placed in the center of the open-field apparatus and left to move freely. Data were individually recorded for each animal during 15 min. Ambulatory walked distance and velocity were measured.

#### Rotarod test

Motor coordination was measured in an accelerating rotarod apparatus (Ugo Basile). Mice were pre-trained during two days at a constant speed, the first day: 4 trials of 1 min at 4 rpm and the second day: 4 trials of 1 min at 8 rpm. On the third day, rotarod was set to accelerate from 4 to 40 rpm over 5 min and mice were tested four times. During accelerating trials, the latency to fall from the rod was measured.

#### Inverted grid test

Muscle strength was examined by placing the mice in the center of a wire grid that was rotated to an inverted position over 10 s and held steadily 50 cm above a padded surface. Latency to fall from the grid was measured (maximum: 300 s, two trials).

#### Social approaching test

Test was conducted in a Plexiglas three-chambered box with openings between the chambers. The first day (habituation), mice were allowed to explore for 10 min the empty box. The next day (test), mice were placed in the center of the box, containing two wire cages placed in opposite corners of the lateral chambers, one empty (“Empty cage”) and the other with an unknown (gender paired) mouse on it (“Occupied cage”). Mice were recorded for 10 min and the time spent in each chamber was measured. Preference index was measured as [time in Occupied cage – time in Empty cage]/[time in Empty cage + time in Occupied cage]*100.

#### Vertical pole

Mice were placed in the top of a 50 cm high vertical 1 cm diameter pole. The pole is mounted on a triangular base stand and placed in the home cage so that mice might prefer to descend to the floor of cage. Mice were trained once to turn around and descend the pole. Then, the animal is placed again in the top of the pole and the recording starts when the animal begins the turning movement. The time to turn completely downward and total time to descend to the floor are measured.

### *In silico* structural analysis

The inserted sequence GLQ was first modeled by replacing residues ^356^YCL^358^ in the crystal structure of human PAH (PDB entry 6HYC) using COOT (44) and selecting rotamers for side chains that diminished steric clashes with nearby elements. Next, we used AlphaFold (45) and ColabFold (46) to predict the structure of the human protein with the wild-type sequence (UniProt P00439) or bearing the 3 amino acid insertion. The analysis of the predicted model was done with COOT. Figures were prepared with PyMOL (Schrödinger, LLC).

### Statistical analysis

The number of individuals used per experiment is indicated in each figure. All results are expressed as mean ± SEM. In all bar plots, individual data points are presented for visualization clarity. Data were analyzed with GraphPad Prism 10. For comparisons of the three experimental groups one-way ANOVA was used. The unpaired *t* test was used to evaluate the significance of differences between two groups. Critical values for significance of *P*<0.05, *P*<0.01 or *P*<0.001 were used throughout the study (significance indicated by *).

## Data availability

The data that support the findings of this study are available upon request from the corresponding author.

## Study approval

All animal studies were approved by the Institutional Ethical Committee for Animal Experimentation (Universidad Autónoma de Madrid, references CEI 963-A026 and CEI-134-2830) and by the Regional Environment Department (Comunidad de Madrid, reference PROEX 194/19).

## Author contributions

L.R.D. conceptualized this study. A.M-P., S.P., A.L-M., C L-R. performed experiments, analysed data and participated in manuscript drafting and preparing of figures. E.M., M.A. and M.C. provided technical support for the experiments. S.R-M. performed in silico structural predictions. B.P., J.J.L., E.R. and L.R.D. provided supervision. E.R. and L.R.D. acquired funding. J.J.L. and L.R.D. drafted and revised the manuscript. All authors revised the final draft and approved submission.

## Acknowledgements

The authors thank Verónica Domínguez and Belén Pintado from CNB-CBM Transgenesis Unit for their help and advice in the generation of the humanized PKU mouse. The help from the Confocal Microscopy (SMOC) and Electron Microscopy core facilities is also acknowledged. This work was funded by Spanish Ministry of Science and Innovation and European Regional Development Fund (grants PID2019-105344RB-I00/AEI/10.13039/501100011033 and PID PID2022-137238OB-I00/AEI/10.13039/501100011033), by Fundación Ramón Areces (XX National Call 2020) and from TaNeDS Europe 2017 (Daichii Sankyo) research grant. Centro de Biología Molecular Severo Ochoa receives an institutional grant from Fundación Ramón Areces.

---

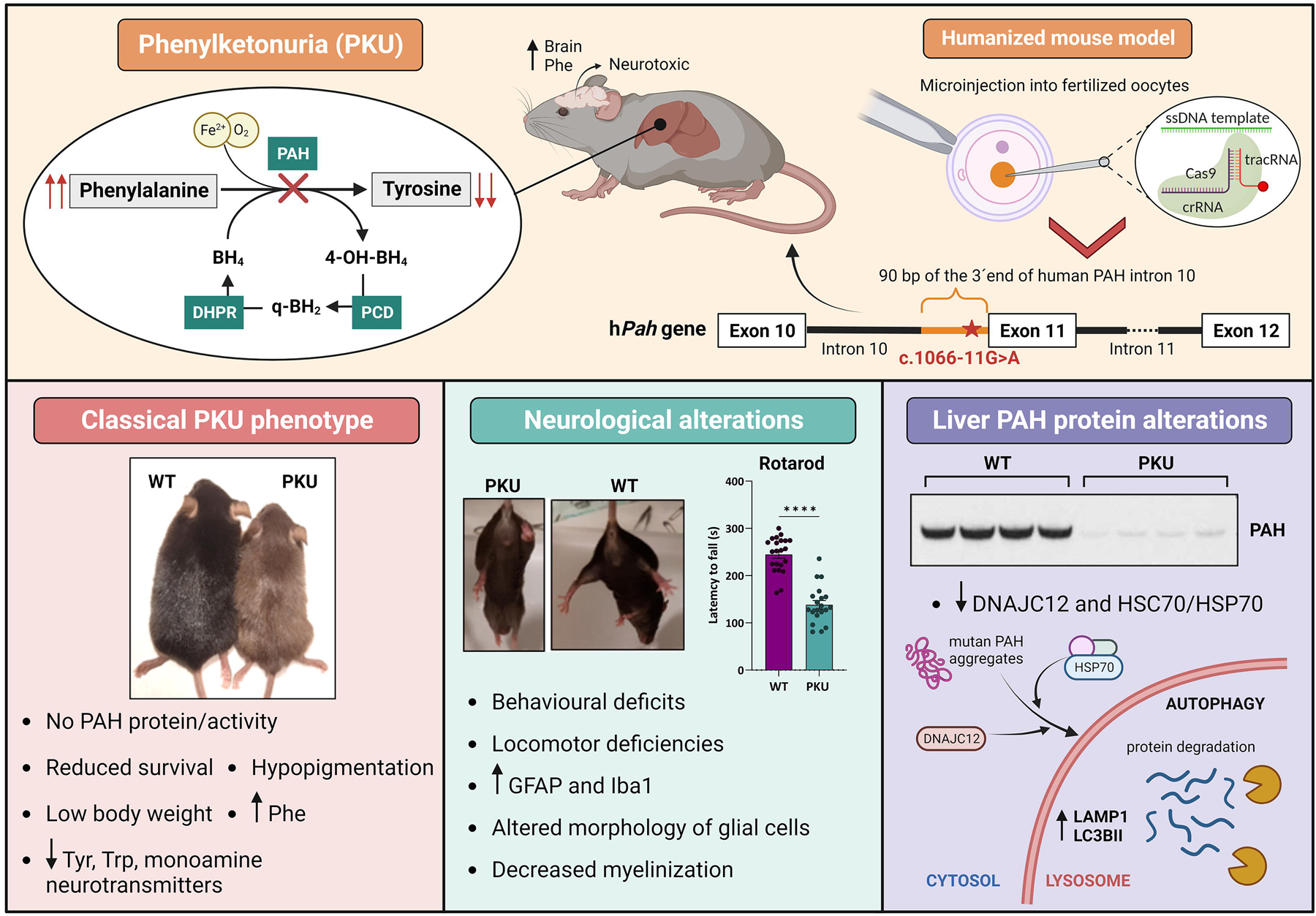

