## Supplementary material for "PAH DEFICIENT PATHOLOGY IN HUMANIZED c.1066-11G>A PHENYLKETONURIA MICE"

| **Supplementary Table 1.** Blood serum concentrations of amino acids, whose levels were found increased/decreases in *Pah* c.1066-11A mice compared to the control WT group. | | | |
|---|---|---|---|
| **Amino acid** | **WT (µM)** | ***Pah* c.1066-11A (µM)** | ***p-value*** |
| **Taurine** | 360.05 ± 17.35 | 284.12 ± 16.17 | 0.0040 |
| **Asparagine** | 41.37 ± 2.31 | 33.62 ± 2.41 | 0.0293 |
| **Proline** | 58.94 ± 2.80 | 49.66 ± 3.25 | 0.0413 |
| **Glycine** | 181.2 ± 10.43 | 143.12 ± 7.17 | 0.0070 |
| **Alanine** | 328.65 ± 15.59 | 249.93 ± 19.37 | 0.0044 |
| **Cystine** | 1.31 ± 0.28 | 10.49 ± 1.27 | 0.0000 |
| **Methionine** | 48.94 ± 1.53 | 41.52 ± 1.69 | 0.0036 |
| **Tyrosine** | 68.20 ± 4.41 | 28.85 ± 1.66 | 0.0000 |
| **Phenylalanine** | 59.26 ± 2.48 | 1885.69 ±71.22 | 0.0000 |
| **Tryptophan** | 55.74 ± 3.78 | 42.15 ± 2.01 | 0.0056 |
| **Lysine** | 245.58 ± 16.57 | 196.64 ± 12.07 | 0.0267 |
| **1-methyl-histidine** | 2.54 ± 0.31 | 1.56 ± 0.27 | 0.0313 |
| **Arginine** | 106.35 ± 6.26 | 68.99 ± 3.95 | 0.0001 |

Concentrations are expressed as mean ± SEM; n=12 WT and n=13 PKU mice.

**Supplementary Table 2.** Brain concentrations of amino acids, whose levels were found increased/decreased in *Pah* c.1066-11A mice compared to the control WT group.

| Amino acid | WT (µM) | *Pah* c.1066-11A (µM) | *p-value* |
|---|---|---|---|
| **Serine** | 145.84 ± 4.51 | 218.47 ± 3.43 | 0.0000 |
| **Glutamine** | 1108.41 ± 46.80 | 822.25 ±17.70 | 0.0001 |
| **Glycine** | 302.5 ± 12.3 | 605.46 ± 15.55 | 0.0000 |
| **Cystathionine** | 3.13 ± 0.19 | 4.37 ± 0.32 | 0.0064 |
| **Isoleucine** | 3.71 ± 0.35 | 2.41 ± 0.21 | 0.0051 |
| **Leucine** | 13.9 ± 0.81 | 9.4 ± 0.39 | 0.0001 |
| **Tyrosine** | 14.58 ± 1.19 | 6.37 ± 0.33 | 0.0000 |
| **Phenylalanine** | 13.62 ± 0.71 | 153.35 ±3.94 | 0.0000 |
| **Lysine** | 42.18 ± 1.2 | 47.55 ± 1.07 | 0.0029 |
| **Histidine** | 12.58 ± 0.51 | 17.31 ± 0.46 | 0.0000 |

Concentrations are expressed as mean ± SEM; n=12 WT and n=13 PKU mice.

**Supplementary Table 3.** Primers used for gDNA and cDNA amplification.

| Name | Sequence 5’-3 | Method |
|---|---|---|
| **PAH 10-11-12 S** | 5'-GGTAACGGAGCCAACATGGTTTACTG-3' | RT-PCR human minigene |
| **PAH 10-11-12 AS** | 5'- AGACTCGAGGGTAGTCTATTATCTGTT-3' | |
| **pcDNA3.1 F** | 5'-ACCCACTGCTTACTGGCTTA-3' | RT-PCR mouse minigene |
| **pcDNA3.1 R** | 5'-GCAACTAGAAGGCACAGTCG-3' | |
| **hPAH exon 10 F** | 5'-ACTGTGGAGTTTGGGCTCTG-3' | RT-PCR HepG2 cells |
| **hPAH exon 12 R** | 5'-ACTGAGAAGGGCCGAGGTAT-3' | |
| **mPAH exon 10 F** | 5’-GGCTCTTGTCATCCTTTGGA-3’ | RT-PCR mice liver |
| **mPAH exon 11 R** | 5’-TACTCCTGGCAGGCTGTCTT-3’ | |
| **hPAH intron 10 F** | 5’-ACCTATGCATGCAGCCTTGT-3’ | PCR HepG2 edited cells (mutation detection) |
| **hPAH intron 11 R** | 5’- AGGGTGGAGAACATGGGAGA-3’ | |
| **mPAH exon 10 F** | 5’-GGCTCTTGTCATCCTTTGGA-3’ | PCR wt allele genotyping |
| **mPAH WT R** | 5’-GCTTCTTCCATCTCCCCATT-3’ | |
| **mPAH exon 10 F** | 5’-GGCTCTTGTCATCCTTTGGA-3’ | PCR mutant allele genotyping |
| **-11G>A human R** | 5’-CAGTATTCCCTGCTGCATCC-3’ | |
| **hPAH ex1-2 F** | 5’-GACTTTGGACAGGAAACAAGC-3’ | qPCR human PAH |
| **hPAH ex2-3 R** | 5’-CATTCTCCTCAAATAAGCGCA-3’ | |
| **hDNAJC12 F** | 5’-CAGTGAAGACGTCAATGCACT-3’ | qPCR human Dnajc12 |
| **hDNAJC12 R** | 5’-GGTTGAAGCCAGCTCCTCT-3’ | |
| **hHSP70 F** | 5’-GATCAACGACGGAGACAAGC-3’ | qPCR human Hsp70 |
| **hHSP70 R** | 5’-GCTGCGAGTCGTTGAAGTAG-3’ | |
| **mPah ex6-7 F** | 5’-TTCTGCAGACTTGTACTGGT-3’ | qPCR mouse PAH |
| **mPah ex7-8 R** | 5’-ACAGATATCAGGTTCAGGTG-3’ | |
| **mDnajc12 F** | 5’-CTCTCCTCGGTTGAGCAAAT-3’ | qPCR mouse Dnajc12 |
| **mDnajc12 R** | 5’-ATGCATCGAAGTTTTCACTG-3’ | |
| **mHsp70 F** | 5’-GCGCTCGAATCCTATGCCTT-3’ | qPCR mouse Hsp70 |
| **mHsp70 R** | 5’-GTGCACGAACTCCTCCTTGT-3’ | |

**Supplementary Table 4.** DNA template, gRNA and top predicted off-targets for gene editing in HepG2 cells.

| | | | | | |
|---|---|---|---|---|---|
| **DNA TEMPLATE** | 5'-CCTATGGGATGCAGCAGGGAATACTGATCCTGATTTAACAGTGATAATAACTTTTCACTTAGGGCCTACAGTACTGCTTA TCAGAGAAGCCAAAGCTTCTCCCCCTGGAGCTGGAGAAGA-3´ | | | | |
| **GUIDE (gRNA)** | 5'-UCUCUGAUAAGCAGUACUGU -3' | | | | |
| **OFF-TARGET SEQUENCE*** | **MM*** | **GENOMIC LOCALIZATION** | **REGION** | **GENE** | **PRIMERS (5'-3')** |
| T**G**T**AG**GA**C**AAGCAGTA CTGTTGG | 4 | Chr8: 118873153-118873175 | INTRON | SAMD12-AS1 | 5'-TGGTGCACTAAAGCAGCCAT-3' 5'-TGCCAAGGCTACGGAAAGTT-3' |
| TCTCT**C**A**C**A**G**GCAGTAC TGTTGG | 3 | Chr8: 118873153-118873175 | UPSTREAM | SCFD2 | 5'-TGGGTTCTGTTTGTTCCACC-3' 5'-AGTGGTTTGATAGGTGGGGG-3' |
| TC**AA**TGA**GG**AGCAGTA CTGTGGG | 4 | Chr21: 39462538-39462560 | EXON | GET1-SH3BGR (IKZF3) | 5'-AGCAGGCACATAAGGGTGTT-3' 5'-GCCCACTGTCCTTAGTGGAT-3' |

*mismatches (MM) shown in bold

**Supplementary Table 5.** DNA template, gRNA and top predicted off-targets for gene editing in mice.

| | | | | | |
|---|---|---|---|---|---|
| **DNA TEMPLATE** | 5′-AGTCCATGCTAAATGTCTCAGTTTTAGGGAATTTGCAAGAGAAGGGGCACAAATGGCCTATGGGATGCAGCAGGGAATACT GATCCTGATTTAACAGTGATAATAACTTTTCACTTGGGGCCTACAGTACTGTTTATCAGACAAGCCAAAGCTCCTGCCCC-3′ | | | | |
| **GUIDE (gRNA)** | 5′-GUUUUAGGGAAUUUGCACUAA-3′ | | | | |
| **OFF-TARGET SEQUENCE*** | **MM*** | **GENOMIC LOCALIZATION** | **REGION** | **GENE** | **PRIMERS (5′-3′)** |
| **T**TT**A**TAGGGAATTTGC ACT**G**TGG | 3 | Chr15: 32795435-32795457 | EXON | Gm32618 | 5′-GGGAACACCAGTTGCTACTAGC-3′<br>5′-GCACCAGAGACTCTCCAGCA-3′ |
| **TA**T**CA**AGGGAATTTGC ACTATGG | 4 | Chr7: 134570396-134570418 | INTRON | Dock1 | 5′-GGTAAAGTCCTCAGGTGGTCA-3′<br>5′-CCATGAGCCAACATTACACC-3′ |
| GTT**A**TA**T**GGA**G**TTTGC ACTAAGG | 3 | Chr7: 89552521-89552543 | EXON | Ccdc81 | 5′-CAGTGCTCTGGCTTCGTGTT-3′<br>5′-CACACACCTGCATCAAGTCA-3′ |
| **TA**T**A**TAGGTAATTTGC ACTAAGG | 4 | Chr10: 120001151-120001173 | INTRON | Irak3 | 5′-GGCAAGAGCAAGAGGAGAAGA-3′<br>5′-GAACCCAGAGCTCTCAGGTCAT-3′ |

*mismatches (MM) shown in bold

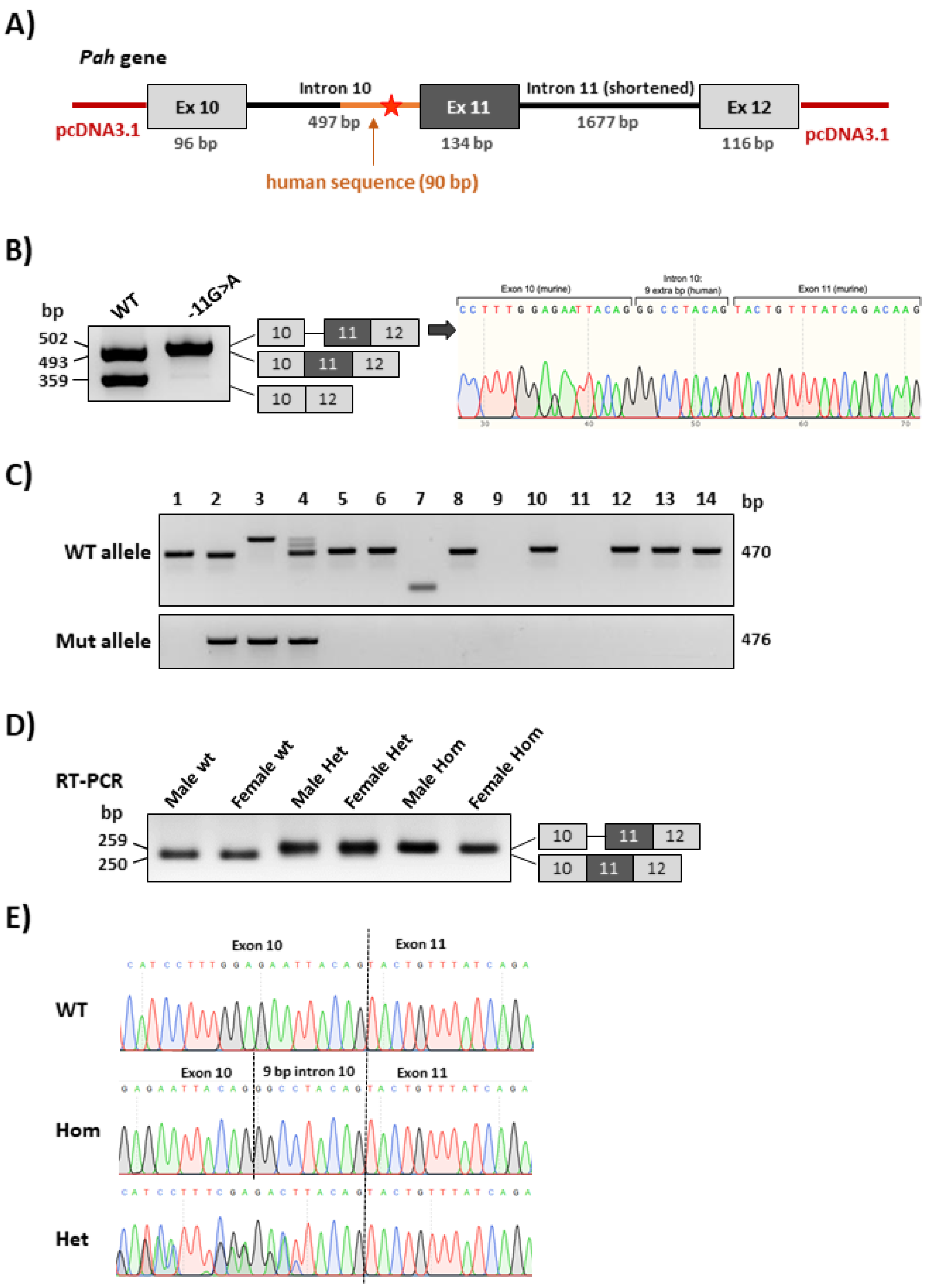


**Supplementary Figure 1. Humanized minigene and *Pah* c.1066-11A (PKU) mice. A)** Schematic representation of minigene construction in the pcDNA3.1 vector. **B)** RT-PCR results after transfection in Hep3B cells of wild-type (WT) and mutant minigenes. Shown on the right is the Sanger sequencing analysis of the PCR product obtained with the mutant minigene. **C)** Gel electrophoresis of PCR genotyping in F0 mice from tail DNA biopsy. **D)** Splicing profile in liver of F5 mice by RT-PCR from WT, heterozygous (Het) and homozygous (Hom) PKU male and female mice. **E)** Sequencing analysis of RT-PCR products from WT, Het and Hom mice.

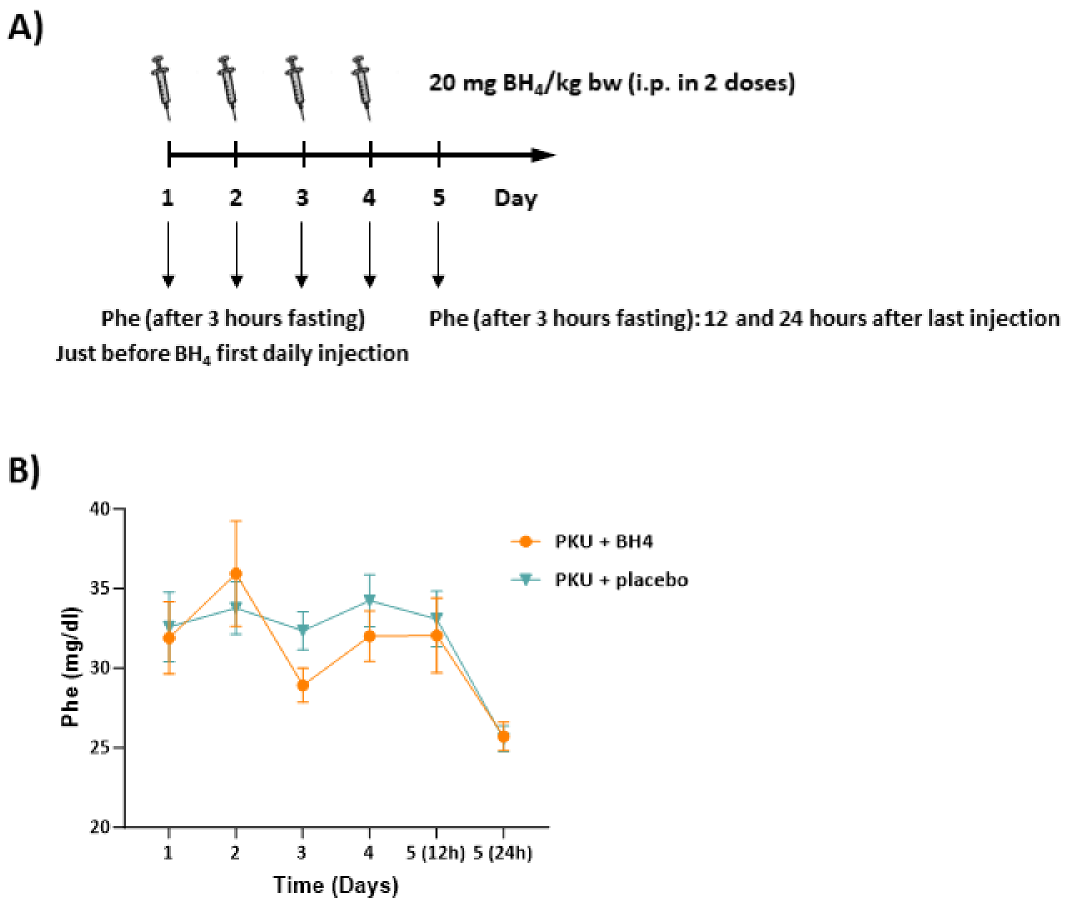


**Supplementary Figure 2. $BH_4$ treatment. A)** Administration protocol of $BH_4$ and **B)** Phe levels after treatment in *Pah* c.1066-11A (PKU) mice. Data are presented as mean of n=5 $BH_4$ treated mice compared to mice treated with placebo.

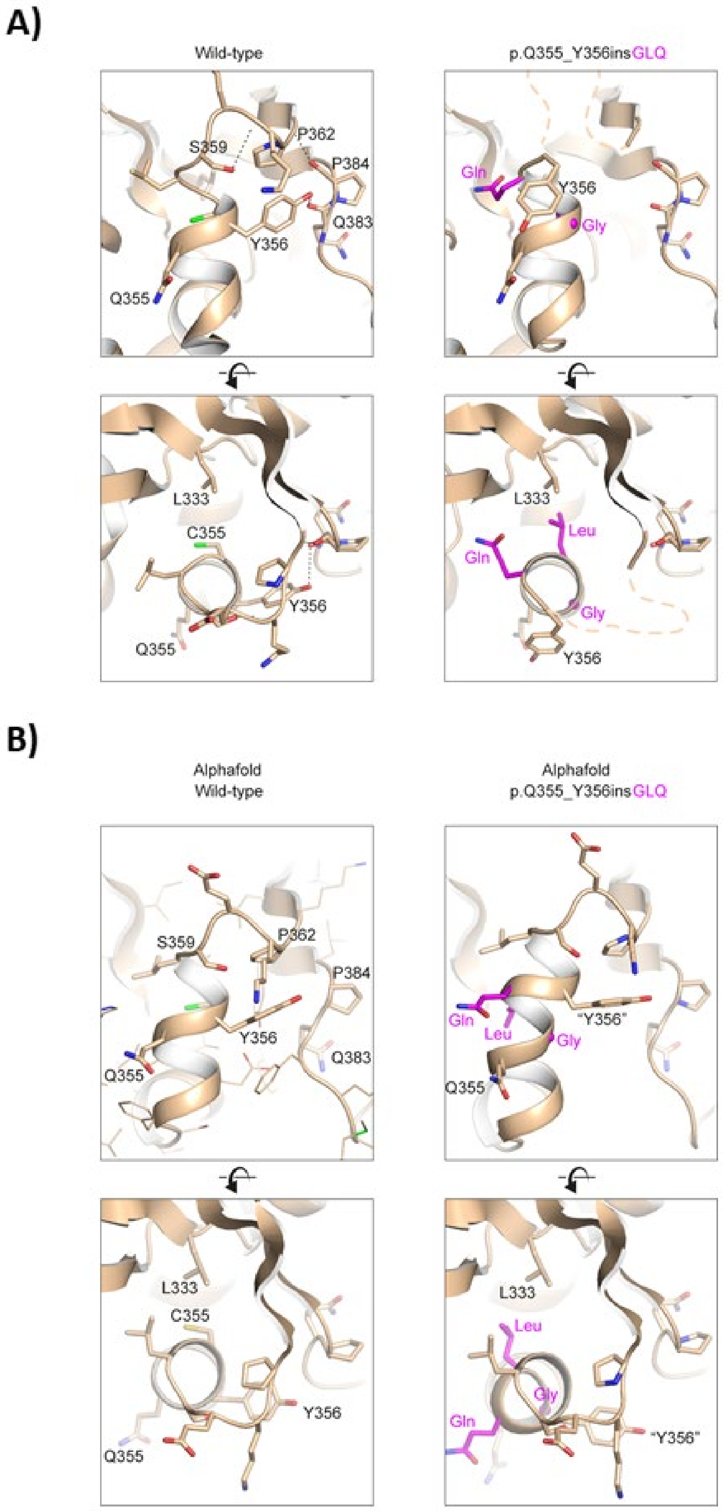


**Supplementary Figure 3.** ***In silico*** **structural analysis of p.Q355_Y356insGLQ. A)** Perpendicular views of the crystal structure of human PAH (PDB entry 6HYC) with relevant residues depicted in sticks and hydrogen bonds as dashed thin lines. To the right, modelling of the pQ355_Y356insGLQ variant, with the three inserted residues depicted in magenta. The thick dashed line indicates that the position of the residues displaced by the insertion is uncertain. **B)** Cartoon representation of the AlphaFold structural predictions for the WT protein (left) and pQ355_Y356insGLQ variant (right).
